# The Tetrameric Structure of Nucleotide-regulated Pyrophosphatase and Its Modulation by Deletion Mutagenesis and Ligand Binding

**DOI:** 10.1101/2020.04.07.028951

**Authors:** Viktor A. Anashkin, Anu Salminen, Victor N. Orlov, Reijo Lahti, Alexander A. Baykov

## Abstract

A quarter of prokaryotic Family II inorganic pyrophosphatases (PPases) contain a regulatory insert comprised of two cystathionine β-synthase (CBS) domains and one DRTGG domain in addition to the two catalytic domains that form canonical Family II PPases. The CBS domain-containing PPases (CBS-PPases) are allosterically activated or inhibited by adenine nucleotides that cooperatively bind to the CBS domains. Here we use chemical cross-linking and analytical ultracentrifugation to show that CBS-PPases from *Desulfitobacterium hafniense* and four other bacterial species are active as 200–250-kDa homotetramers, which seems unprecedented among the four PPase families. The tetrameric structure is stabilized by Co^2+^, the essential cofactor, pyrophosphate, the substrate, and adenine nucleotides, including diadenosine tetraphosphate. The deletion variants of *dh*PPase containing only catalytic or regulatory domains are dimeric. Co^2+^ depletion by incubation with EDTA converts CBS-PPase into inactive tetrameric and dimeric forms. Dissociation of tetrameric CBS-PPase and its catalytic part by dilution renders them inactive. The structure of CBS-PPase tetramer was modelled from the structures of dimeric catalytic and regulatory parts. These findings signify the role of the unique oligomeric structure of CBS-PPase in its multifaced regulation.

## 1. Introduction

Pyrophosphate (PP_i_), a byproduct of numerous biosynthetic reactions [1], re-enters metabolic pathways after being converted to phosphate via the action of specific enzymes—pyrophosphatases (PPases; EC 3.6.1.1). Soluble PPases belong to three non-homologous families, I, II, and III [2–4]. Family I PPases are found in all kingdoms of life, whereas Family II PPases and the less common Family III PPases are found in prokaryotes. With rare exceptions [5], PPases of Families I and III are single-domain proteins.

Family II PPases have a more variable domain organization and are divided into two subfamilies. In the “canonical” subfamily, which has been well characterized both structurally and mechanistically [6], each subunit of 33–34 kDa is formed by two catalytic domains, DHH and DHHA2, connected by a flexible linker, and the active site is located between the domains [7, 8] (Fig. 1A). These PPases form stable homodimers in physiological conditions [9], via association of the DHH domains [7, 8], and are Mn^2+^- or Co^2+^-metalloproteins that additionally bind several Mg^2+^ ions less tightly [9]. The metal ions stabilize the oligomeric structure of PPases [9] and are the key to catalysis; in the latter function they neutralize and position the substrate PP_i_ and convert the nucleophilic water molecule to a highly reactive hydroxide ion [10].

**Fig. 1.**
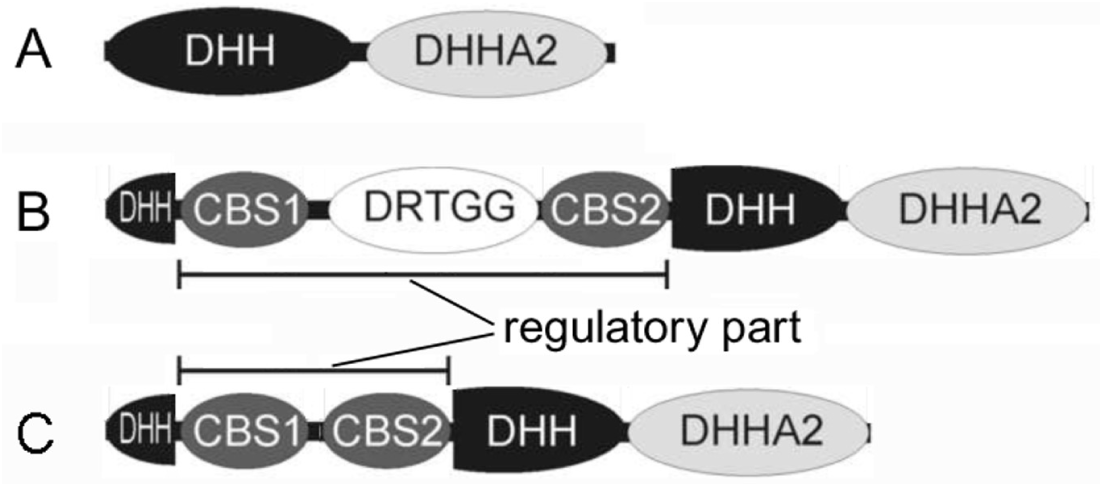
Domain topologies of canonical Family II PPase (A) and CBS-PPase (B,C). DHH and DHHA2 are catalytic domains; CBS1, CBS2 and DRTGG domains form a regulatory part within the DHH domain.

A quarter of Family II PPases, belonging to another subfamily (named “CBS-PPases”), contain a regulatory insert of ∼250 residues comprised of two β-cystathionine synthase (CBS) domains and, frequently, one DRTGG domain within the DHH domain (Fig. 1B,C) [11,12]. CBS-PPases are unique among different PPases in being differentially regulated (activated or inhibited) by adenine mono- and dinucleotides that bind to the CBS domains [11,13,14]. This property is thought to enhance the control of PP_i_ level under stress conditions and, hence, the viability of the bacterial host [14]. The regulating nucleotides bind to the enzyme in a positively co-operative manner [13], a phenomenon associated with the oligomeric structure of CBS-PPase.

The three-dimensional structure of the full-size CBS-PPase remains to be determined. Tuominen et al. [12] have reported on the crystal structure of the isolated regulatory part (residues 66–306) of *Clostridium perfringens* CBS-PPase. The regulatory part is dimerized by forming CBS-CBS′ and DRTGG-DRTGG′ contacts (prime sign refers to the other subunit) and retains the ability to bind adenine nucleotides [12], which means that its native structure is well preserved in the absence of the catalytic domains. Accordingly, deletion of the regulatory domains did not inactivate CBS-PPase but abolished nucleotide effects on activity [13]. The catalytic domains of CBS-PPases should have a fold very similar to that of the canonical PPases, based on high sequence identity. Based on this information, a model of full-size CBS-PPase with a complete set of domains was suggested, which assumed its dimeric organization [12].

Here we provide experimental evidence that, contrary to the above prediction, bacterial CBS-PPases form homotetramers. Most of the experiments described in this manuscript were performed with *Desulfitobacterium hafniense* CBS-PPase (*dh*PPase), which contains both CBS and DRTGG domains in its regulatory part [13]. However, critical controls run with four other CBS-PPases, including those lacking DRTGG domain, indicated that they are also tetrameric proteins. We have also determined the contributions of the catalytic and regulatory parts to tetramer stability and its dependence on substrate and adenine nucleotide binding to the catalytic and regulatory sites, respectively.

### 2. Materials and methods

#### 2.1 Enzymes

*D. hafniense* CBS-PPase (*dh*PPase), its separate catalytic part (*dh*PPaseΔCDC), and the CBS-PPases of *Clostridium perfringens* (*cp*PPase), *Clostridium novyi* (*cn*PPase), *Eggerthella lenta* CBS-PPase (*el*PPase), and *Ethanoligenens harbinense* CBS-PPase (*eh*PPase) were produced and purified as described elsewhere [12, 13, 15]. The last two enzymes (*el*PPase and *eh*PPase) lack the DRTGG domain in their structures. The gene for a separate regulatory part of *dh*PPase (*dh*CDC, residues 63– 306 of *dh*PPase) was produced using overlap-extension PCR with Phusion DNA polymerase and the primers listed in Table S1. The mutant gene was ligated into the NdeI and XhoI restriction sites of the pET-42b plasmid (Novagen) and transformed into *Escherichia coli* DH5α cells. The construct was confirmed by sequencing.

All CBS-PPases and their deletion variants were produced in *Escherichia coli* BL21 cells. Collected cells with produced *dh*CDC were suspended in lysis buffer (25 mM Tris-HCl, 10 mM MgCl_2_, pH 7.3) and disrupted with 2–3 cycles of freezing in liquid nitrogen followed by thawing in a water bath at room temperature in the presence of 0.2 mg/ml lysozyme. Genomic DNA was destroyed by DNAse I (Roche, Germany). The *dh*CDC was isolated from the cell extract by ion-exchange chromatography in the lysis buffer with a linear gradient of 0–0.3 M NaCl followed by gel filtration in 50 mM Hepes/KOH, pH 7.5, containing 2 mM MgCl_2_ and 25 mM KCl. *dh*CDC was identified in eluate by SDS-PAGE in the Laemmli system [16]. The *dh*CDC-containing fractions obtained after gel filtration were divided into aliquots, frozen, and stored at -50 °C. The protein was electrophoretically homogeneous.

Protein concentrations were determined with a Nanodrop spectrophotometer (Thermo Scientific) using subunit molecular mass/*A*^0.1%^_280_ of 60.4 kDa/0.478 for *dh*PPase, 34.5.kDa/0.419 for *dh*PPaseΔCDC, 25.9 kDa/0.541 for *dh*CDC, 63.6 kDa/0.548 for *cn*PPase, 60.8 kDa/0.426 for *cp*PPase, 52.5 kDa/0.493 for *el*PPase, and 49.8 kDa/0.478 for *eh*PPase, as calculated from their amino acid compositions with ProtParam. Molar concentrations of PPases are reported below in terms of the monomer.

### 2.2. Biochemical methods

The initial rates of PP_i_ hydrolysis were measured using a continuous P_i_ assay [17] at a sensitivity of 2–50 μM P_i_ per recorder scale. The assay medium contained 140 μM PP_i_ (yielding 50 μM MgPP_i_ complex), 0.1 M Tes-KOH, pH 7.2, 5 mM MgCl_2_ and 0.1 mM CoCl_2_, except where specified. The reaction was initiated by adding enzyme and continued for 2–3 min at 25 °C. The dead time of the phosphate analyzer (i.e., the time from feeding the sample into the analyzer to mixing it with the quenching acidic molybdate solution) was 10-20 s in the standard mode. To increase the precision of the initial velocity estimation for reactions yielding non-linear P_i_ production traces, the dead time was decreased to 1 s by an appropriate modification of the inlet system [18]. The rate values obtained in replicate measurements usually agreed within 5–10%.

Cross-linking of CBS-PPase with glutaraldehyde was performed for 15 min at 25 °C in 25 mM HEPES-KOH buffer, pH 7.5. The reaction was terminated by adding 1/10 volume of 1 M Tris/HCl, pH 7.5. Cross-linked samples were diluted to 0.3 mg/ml protein and separated by electrophoresis in 4–12% gradient polyacrylamide gels, using identical protein loads (9 µg per lane), on a Mini Protein II apparatus (Bio-Rad) in the Laemmli system [16]. Prestained PAGE rulers (Thermo Fisher Scientific) were used as molecular mass standards. The gels were fixed for 20 min with 10% acetic acid/25% isopropanol, stained for 40 min with 0.15% Coomassie G-250 and 0.024% CuSO_4_ in 10% acetic acid/30% ethanol, and destained with 5% acetic acid/15% ethanol.

Analytical ultracentrifugation was performed at 25 °C in a Spinco E instrument (Beckman Instruments) equipped with a computerized data collection unit, with scanning at 280 nm. Samples containing 10–20 µM proteins in the isolation buffer were preincubated for 6 h at 25 °C, unless otherwise indicated. MgCl_2_ and CoCl_2_ were not added to the samples containing EDTA. The sedimentation velocity was measured at 60,000 rpm, and sedimentation coefficients (*s*_20,w_) and molecular masses were estimated with the program SedFit [19].

### 2.3. Data analysis

Nonlinear least-squares fittings were performed using the program SCIENTIST (Micromath), which allows use of differential and implicit equations. Equations 1 and 2, derived from Scheme 1, describe time-courses of activity (*A*) resulting from tetramer (T) dissociation into dimers (D). At *d*α_T_/*dt* = 0, these equations describe the equilibrium activity as a function of enzyme concentration. *A*_T_ and *A*_D_ are specific activities of tetramer and dimer, respectively, α_T_ is the fraction of enzyme in tetrameric form at time *t*, [E]_t_ is total enzyme concentration, expressed in monomers, *k*_a_ and *k*_d_ are the apparent rate constants for tetramer formation and breakdown, respectively.

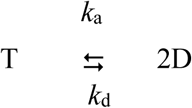

**Scheme 1**. Reversible dissociation of tetrameric enzyme into dimers.

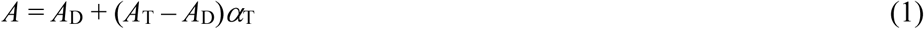

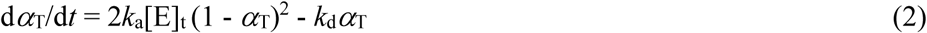

The time-courses of PP_i_ hydrolysis by *dh*PPaseΔCDC, accompanied by first-order inactivation (dissociation) of the enzyme, were fit to Equation 3, where [P] is phosphate concentration, *k*_i_ is the rate constant for enzyme inactivation, *v*_0_ is the initial rate of product formation, *t* is time, and *a* is the background signal of the instrument.

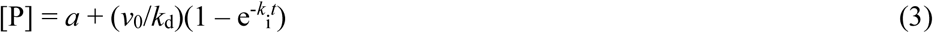

### 2.4. Homology modelling

The full-length *cp*PPase model was constructed using *cp*PPase amino acid sequence and the structures of canonical Family II PPase of *Bacillus subtilis* and AMP-complexed *cp*CBS (PDB IDs: 2HAW and 3L31) [12] as templates. These dimeric structures were manually arranged, using UCSF Chimera v. 1.11.2 [20], into a preliminary tetrameric form that was used as the input for Modeller v. 9.23 [21] operated in “automodel” mode with restrained known contacts within dimers.

## 3. Results

### 3.1. Cross-linking of CBS-PPase

Cross-linking experiments provided strong evidence for the tetrameric structure of CBS-PPase. Full-size *dh*PPase was reacted with glutaraldehyde, using a reaction time and cross-linker concentration that allowed only partial modification, in order to minimize intermolecular cross-linking. To estimate its extent, the reaction was performed at various protein concentrations, keeping in mind that only the intermolecular but not intramolecular reaction should depend on the concentration. The reaction products were separated by SDS-PAGE (Fig. 2A). Dimer and tetramer were the major cross-linked species, but for an unknown reason each migrated as two–three close bands. The intensity of the bands attributable to tetramer did not depend on protein concentration, ruling out intermolecular cross-linking.

**Fig. 2.**
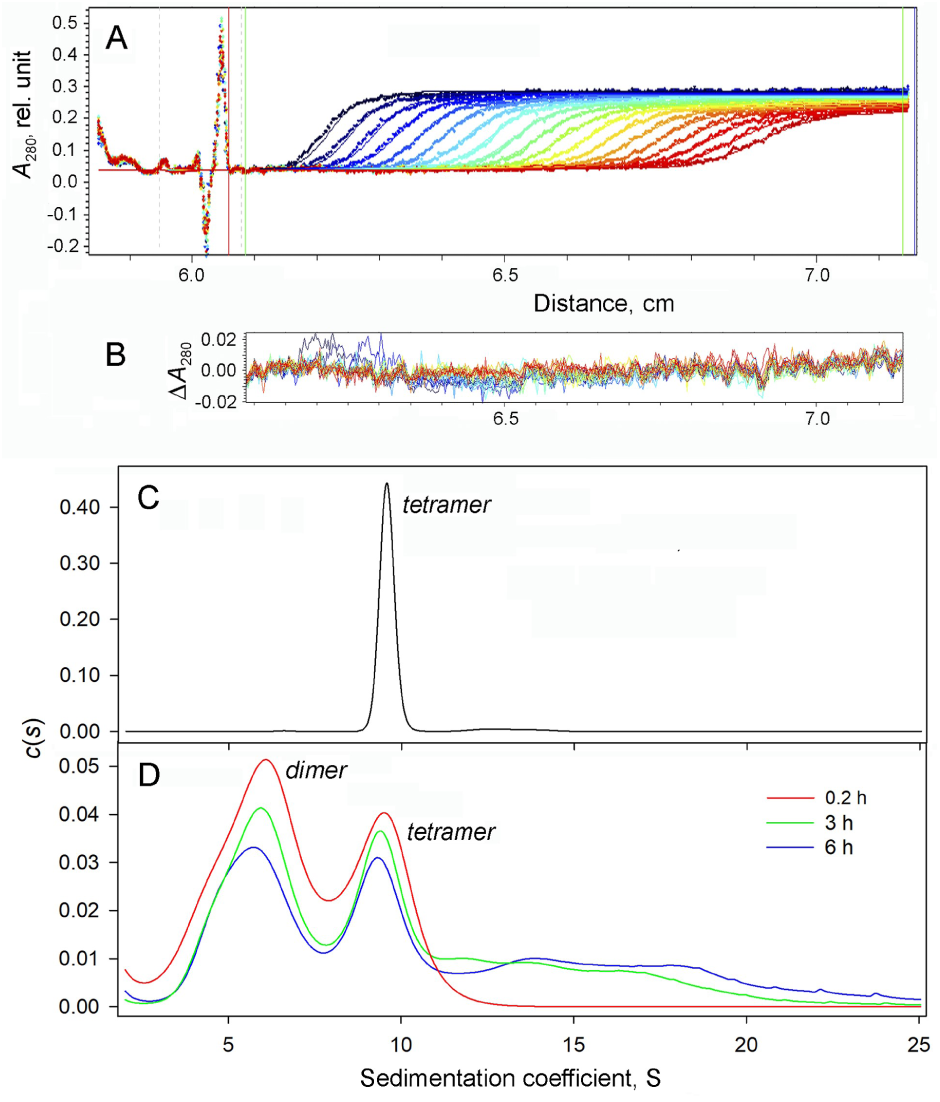
PAGE of *dh*PPase and its deletion variants crosslinked for 15 min with glutaraldehyde. Protein identity and concentration, and glutaraldehyde (GA) concentration for each lane are shown above the gels, as detailed in panel A. The central lane with indicated zero glutaraldehyde concentration refers to non-crosslinked protein in each panel, the left-side lane shows molecular mass markers with the indicated masses in kDa.

Both deletion variants of *dh*PPase produced only dimers upon cross-linking (Fig. 2B,C). Cross-linked *dh*CDC, but not *dh*PPaseΔCDC dimer also migrated as two close bands in SDS-PAGE (Fig. 2C), indicating that the regulatory part was responsible for the unusual behavior of full-size *dh*PPase in SDS-PAGE (Fig. 2A). Cross-linking of an equimolar mixture of *dh*CDC and *dh*PPaseΔCDC (20 µM each) yielded only homodimers of each separate protein, but not their heterodimer (Fig. S1).

Similar results indicating tetrameric structure were obtained with *cn*PPase and *cp*PPase (Fig. S2). With *eh*PPase and *el*PPase, which lack DRTGG domain in their structures, reaction with glutaraldehyde under a variety of conditions (reaction time and glutaraldehyde concentration) yielded high-mass aggregates that did not enter polyacrylamide gel. Dimethyl suberimidate, at a concentration of up to 20 mM (2 h reaction time), generated a small amount of dimers with *eh*PPase (Fig. S3) and no cross-linked products with *el*PPase (data not shown).

### 3.2. Sedimentation velocity analysis of the oligomeric structure of CBS-PPase

Sedimentation velocity measurements were performed with all full-size CBS-PPases and two deletion variants without regulatory or catalytic parts. Figure 3 shows representative data for the full-size *dh*PPase, and Table 1 summarizes the final results for three *dh*PPase forms and all other CBS-PPases, as generated by the SedFit program [19]. Sedimentograms revealed tetramers as the major species in all full-size CBS-PPases. The identification of the major species with s_20,w_ of 9.3-10.2 S in *dh*PPase, *cn*PPase, and *cp*PPase as tetramers was based on the results of the cross-linking experiments and was consistent with the s_20,w_ values reported for proteins of known oligomeric composition [22]. In contrast, all deletion variants, representing the catalytic or regulatory part of CBS-PPases, were predominantly dimeric. Species larger than tetramers were observed in minor amounts in the full-size PPases but were absent in their deletion variants.

**Table 1.** Sedimentation velocity data. Values separated by slash refer to different oligomeric forms. Samples contained 10 µM full-length CBS-PPases or 20 µM their truncated variants. AMP (200 µM) and Ap_4_A (50 µM) were added to protein samples immediately before the sedimentation run; 1 mM EDTA was added immediately before the run (the time of incubation with EDTA during rotor acceleration was roughly estimated as 0.2 h), 3 h prior to the run, or 6 h prior to the run.

| Enzyme | Ligand/additive | $s_{20,w}^a$, S | Fraction distribution<sup>a</sup>, % | Estimated mass of the major species<sup>b</sup>, kDa |
| --- | --- | --- | --- | --- |
| *dhPPase* | None | 9.6 / 13.1 | 96 / 3.7 | 203–234 (T, 241) |
|  | EDTA (0.2 h) | 6.0 / 9.5 | 61 / 38 | 103–115 (D, 121) |
|  | EDTA (3 h) | 5.8 / 9.4 / 12.0 / 14.0 / 16.6 | 45 / 31 / 8 / 9 / 7 | 98–110 |
|  | EDTA (6 h) | 5.5 / 9.3 / 13.8 / 17.8 / 21.4 | 40 / 29 / 16 / 9 / 5 | 90–101 |
|  | AMP | 5.6 / 9.6 / 12.2 | 10 / 84 / 6 | 203–234 |
|  | Ap<sub>4</sub>A | 6.3 / 9.9 / 14.3 | 5 / 91 / 4 | 217–245 |
| *dhPPase*ΔCDC | None | 4.0 | 98 | 56–63 (D, 69) |
|  | EDTA (0.2 h) | 3.1 / 5.8 | 93 / 6 | 38–43 (M, 34.5) |
| *dhCDC* | None | 4.0 | 99 | 56–63 (D, 52) |
|  | EDTA (0.2 h) | 4.0 | 99 | 56–63 |
| *cpPPase* | None | 5.5 / 10.2 / 12.5 | 12 / 80 / 6 | 227–254 (T, 243) |
| *cnPPase* | None | 9.3 / 13.1 | 95 / 4 | 198–222 (T, 254) |
| *ehPPase* | None | 8.2 / 12.0 | 93 / 7 | 173–194 (T, 199) |
| *elPPase* | None | 8.5 / 12.0 | 91 / 8 | 164–184 (T, 210) |
| *sgPPase* (Mn)<sup>c</sup> | None | 3.8 |  | 68 (D, 67) |
| *sgPPase* (Mn)<sup>c</sup> | EDTA | 3.1 |  | 32 (M, 33.6) |

<sup>a</sup> Values separated by slash refer to different oligomeric forms.
<sup>b</sup> Masses were estimated assuming the range of the frictional ratio, $f/f_0$ , of 1.25–1.35, typical for globular proteins [20]. The theoretical masses of the closest oligomeric forms (tetramer, dimer or monomer) are shown in parentheses and marked as T, D or M, respectively.
<sup>c</sup> From Parfenyev et al. [9].

**Fig. 3.**
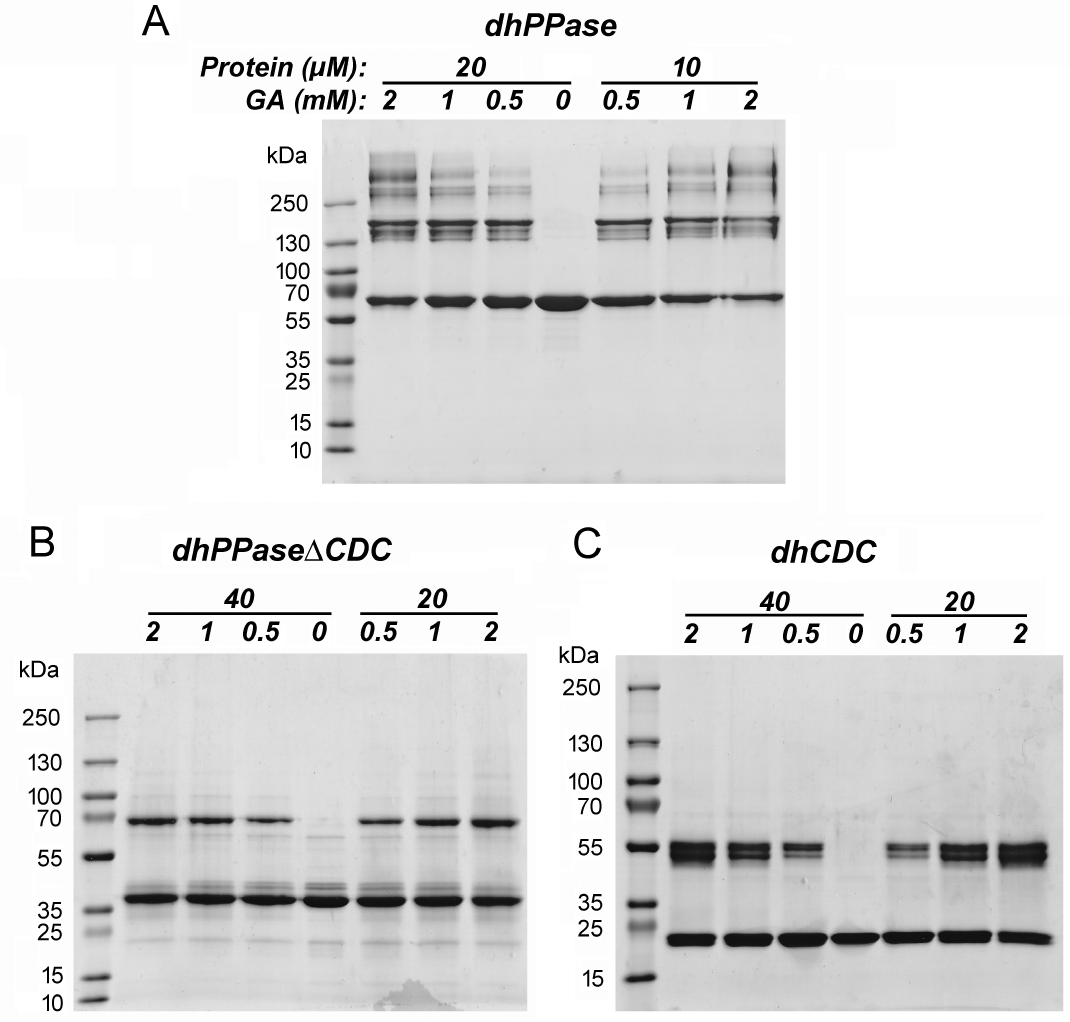
Sedimentation data for *dh*PPase. (A) Raw data for freshly isolated *dh*PPase in the absence of EDTA and nucleotides. The sample contained 20 µM *dh*PPase in the isolation buffer. Red horizontal line is the baseline, red vertical line indicates the position of the meniscus, green vertical lines are the boundaries of the area used in the calculations. (B) The distribution of residuals of the fit in panel A, as generated with SedFit. (C) The distribution of sedimentation coefficients generated with SedFit from the data for untreated *dh*PPase in panel A. (D) Same for *dh*PPase subjected to a 0.2-, 3- or 6-h preincubation with 1 mM EDTA before the sedimentation run. EDTA (1 mM) was added immediately before the run (the preincubation time was roughly estimated as 0.2 h), 3 h prior to the run, or 6 h prior to the run.

Preincubation of protein samples for 0.2 h with 1 mM EDTA before sedimentation induced partial dissociation of tetrameric *dh*PPase into dimers and monomers (Fig. 3D). The *c*(*s*) distribution was characteristic of a mixture of rapidly interconverting tetramers, dimers, and monomers – the tetramer peak was shifted to lower *s* values, and all peaks were asymmetrical, wide and not well separated. A longer preincubation with EDTA increased the fraction of monomers, as indicated by a further shift of the peaks to lower *s* values and appearance of a clear low-mass shoulder and stimulated, in addition, conversion of dimeric *dh*PPase into larger enzyme species (Fig. 3E). Dimeric *dh*PPaseΔCDC was nearly completely dissociated into monomers in the presence of EDTA (Table 1), whereas EDTA-treated *dh*CDC retained its dimeric structure (Table 1). The allosteric inhibitor AMP and activator 5’,5-P1,P4-diadenosine tetraphosphate (Ap_4_A) exhibited minor effects, if any, on the quaternary structure of *dh*PPase (Table 1).

### 3.3. Dilution-induced dissociation of CBS-PPase

Changes in the oligomeric state of the catalytically active *dh*PPase forms could be conveniently monitored by activity measurements, because activities of the associated and dissociated forms were different and they converted into each other slowly on the time scale of the enzyme assay, like with canonical Family II PPases [9]. Dilution of *dh*PPase into the same medium induced its time-dependent inactivation until a constant non-zero activity value was attained (Fig. 4A). Omission of Mg^2+^ from the medium did not change the inactivation time-course (data not shown). EDTA, which depleted the enzyme from both Co^2+^ and Mg^2+^, stimulated the inactivation and decreased the final activity level to nearly zero. The *dh*PPaseΔCDC variant was inactivated much faster upon dilution, and the inactivation was only slightly stimulated by EDTA (Fig. 4B). The final level of activity attained after equilibration of the diluted enzyme increased with increasing enzyme concentration in the absence of EDTA but remained at a nearly zero level in the EDTA-containing medium (Fig. 5).

**Fig. 4.**
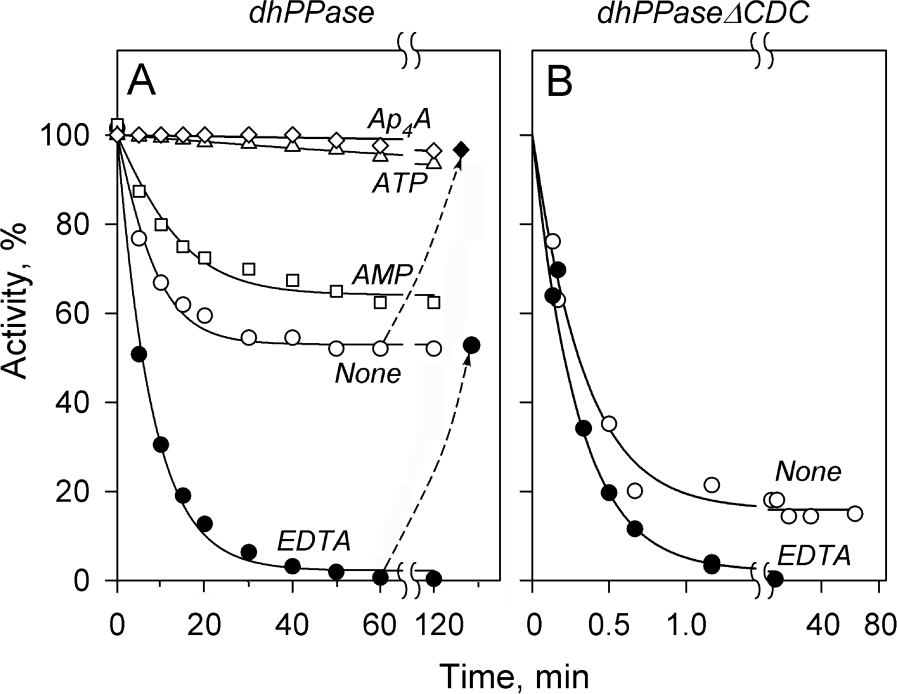
Time-courses of *dh*PPase (A) and *dh*PPaseΔCDC (B) activity upon dilution at 25 °C. Stock enzyme solution (70 μM *dh*PPase or 100 μM *dh*PPaseΔCDC, 0.1 M Mops/KOH, pH 7.2, 2 mM MgCl_2_ and 0.1 mM CoCl_2_) was diluted to 0.8 μM (*dh*PPase) or 2 μM (*dh*PPaseΔCDC) into the same buffer containing no other additions (○), 0.1 mM AMP (□), 0.1 mM ATP (Δ), 5 μM Ap_4_A (◊) or 1 mM EDTA (•). Aliquots were withdrawn in time, and activity was measured in the standard assay medium. The lines represent the best fits for Eqns 1 and 2 and were created using parameter values found in Table 2. Activity extrapolated to zero time (typically 290 s^-1^) was taken as 100 % for each inactivation curve. The dashed lines ending with symbols show reactivation achieved in 60–65 min upon addition of 5 µM Ap_4_A to the sample incubated for 60 min without additions (---♦; the final activity value was corrected for activation due to Ap_4_A carryover to the activity assay medium), or upon EDTA removal by centrifugal ultrafiltration from the sample incubated for 60 min with EDTA (---•).

**Fig. 5.**
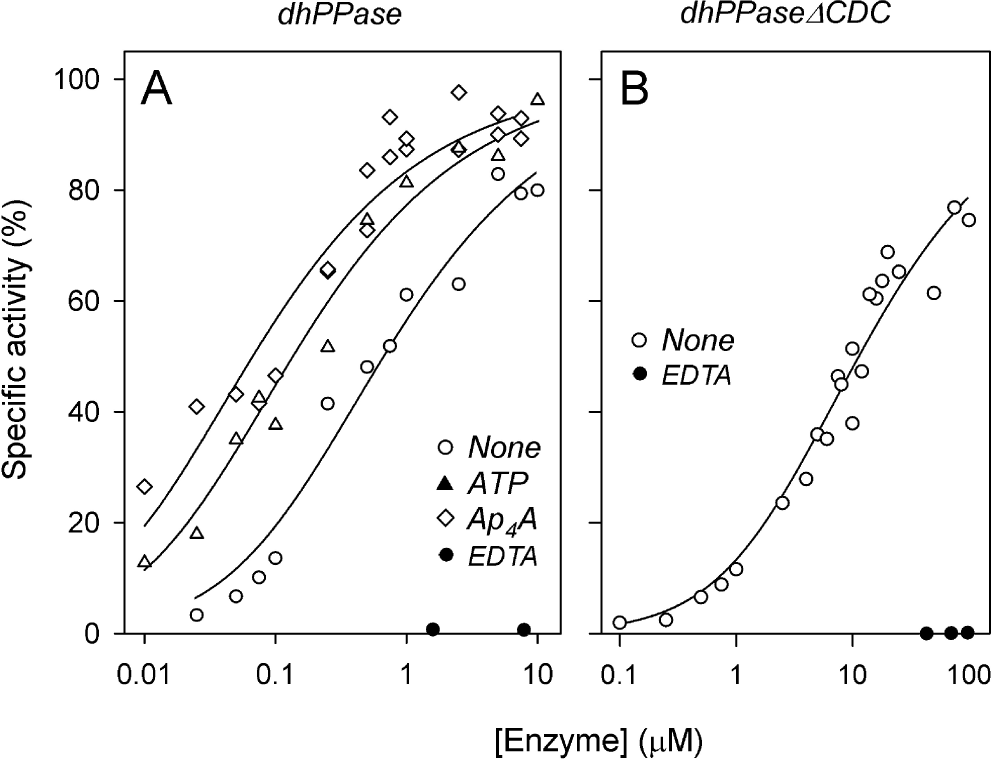
Specific activities of *dh*PPase (A) and *dh*PPaseΔCDC (B) pre-equilibrated at 25 °C at different protein concentrations. The incubation medium contained 0.1 M Mops-KOH buffer, pH 7.2, 2 mM MgCl_2_, 0.1 mM CoCl_2_, and one of the following additions, where indicated: 100 µM ATP, 5 µM Ap_4_A, 1 mM EDTA (symbols are defined on the panels). The metal salts were omitted in the incubations with EDTA. After a 2-h equilibration, aliquots were withdrawn and enzymatic activity was measured in the standard assay medium. The lines represent the best fits of Eqns 1 and 2 with *dα*_D_/*dt* = 0, using the parameter values found in Table 2. Activity extrapolated to infinite enzyme concentration was taken as 100 % for each fitted curve.

The time-courses in Fig. 4 could be analyzed in terms of a model assuming reversible dissociation of CBS-PPase into two equal part (dimers with *dh*PPase and monomers with *dh*PPaseΔCDC). The data in Fig. 4 were fit to Eqns 1 and 2 with dαd/d*t* set to zero, yielding parameter values summarized in Table 2. Lowering the temperature to 0 °C slightly decreased *k*_d_ and *k*_a_ for both *dh*PPase and *dh*PPaseΔCDC with no or minor effect on *K*_d_ (Table 2).

**Table 2.** Parameter values for oligomerization equilibria in *dh*PPase and canonical Family II *sg*PPase derived from activity measurements. Values of *k*_d_, *k*_a_ and *K*_d_ were estimated with Eqns 1 and 2 from inactivation time-courses (Fig. 4) and/or from equilibrium activity versus enzyme concentration profiles (Fig. 5). Ligand concentrations were as follows: AMP, ADP and ATP, 100 µM; Ap_4_A, 5 μM; EDTA, 1 mM. Incubations were performed at 25 °C in 0.1 M Mops/KOH buffer, pH 7.2, containing 2 mM MgCl_2_ and 0.1 mM CoCl_2_, except where otherwise noted.

| Enzyme | Ligand/Additive | $k_d$, h<sup>-1</sup> | $k_a$, μM<sup>-1</sup>h<sup>-1</sup> | $K_d$, μM |
| --- | --- | --- | --- | --- |
| *dh*PPase | None | 2.7 ± 0.2 | 4.0 ± 0.4 | 0.7 ± 0.2 |
| | None (0 °C) | 1.80 ± 0.1 | 2.4 ± 0.5 | 0.7 ± 0.2 |
| | AMP | 1.3 ± 0.2 | 2.5 ± 0.4 | 0.52 ± 0.03 |
| | ADP | 1.14 ± 0.06 | 4.4 ± 0.3 | 0.26 ± 0.01 |
| | ATP | | | 0.14 ± 0.03 |
| | Ap<sub>4</sub>A | | | 0.07 ± 0.01 |
| | EDTA | 7.3 ± 0.4 | | |
| | EDTA + AMP | 6.9 ± 0.3 | | |
| | EDTA + ATP | 7.0 ± 0.3 | | |
| | EDTA + Ap<sub>4</sub>A | 6.9 ± 0.4 | | |
| *dh*PPaseΔCDC | None | 144 ± 6 | 9 ± 3 | 17 ± 5 |
| | None (0 °C) | 70 ± 10 | 7 ± 2 | 10 ± 2 |
| | EDTA | 210 ± 20 | | |
| *sg*PPase (Mn)<sup>a</sup> | None | <1 | 1210 ± 60 | <0.001 |
| | EDTA | 0.27 ± 0.01 | 0.0029 | 93 ± 10 |

<sup>a</sup> From Parfenyev et al. [9]; the parameter values shown refer to enzyme that contained Mn<sup>2+</sup> in the high-affinity site.

The following experiment indicated that the dilution-induced inactivation of *dh*PPase in the presence of EDTA was completely reversible. EDTA-treated *dh*PPase with a nearly zero activity (Fig. 4A) was stripped of EDTA and reconstituted with Co^2+^ by three cycles of a 10-fold concentration on a Centricon centrifugal filter and dilution with the buffer containing 100 μM CoCl_2_ and 2 mM MgCl_2_ instead of EDTA. This procedure reactivated *dh*PPase by 54% (i. e., to the level observed in Fig. 4A) after prolonged incubation in the absence of ligands. The easy reversibility of *dh*PPase inactivation is consistent with EDTA-induced Co^2+^ depletion and/or enzyme dissociation into inactive parts upon dilution.

### 3.4. Stabilization of oligomeric structure by regulatory and active site ligands

The dilution-induced inactivation (tetramer dissociation) of *dh*PPase was less pronounced in the presence of AMP and, especially, ATP or Ap_4_A (Fig. 4A). Accordingly, the dependences of the specific activity on enzyme concentration in the equilibrated system were shifted to lower concentrations in the presence of the adenosine phosphates (Fig. 5A). Furthermore, the activity of *dh*PPase half-inactivated by incubation in the diluted state in the absence of ligands was completely restored upon a 70-min incubation with 5 µM Ap_4_A (Fig. 4A). Of note, the nucleotides were ineffective with the full-size enzyme in the presence of EDTA or with *dh*PPaseΔCDC (data not shown). In terms of Scheme 1, nucleotide effects on *K*_d_, at least for AMP and ADP, resulted mainly from a change in *k*_d_ and increased with the size of the nucleotide (Table 2).

The effect of PP_i_ on oligomer stability could not be tested in the same way because PP_i_ is rapidly consumed by CBS-PPase. To overcome this difficulty, we estimated rates of enzyme dissociation from the time-courses of *dh*PPaseΔCDC-catalyzed PP_i_ hydrolysis at different levels of enzyme saturation with substrate (MgPP_i_ complex). Two MgPP_i_ concentrations were used — 5 and 100 µM, corresponding to 56 and 95% saturation, respectively, based on the *K*_m_ value of 4 µM [13]. The choice of *dh*PPaseΔCDC for this experiment is explained by its propensity to dissociate on the time scale of the enzyme assay compared with the full-size enzyme. At the low concentration of the enzyme used in the activity assay (10^−9^ M), dimer dissociation and, hence, enzyme inactivation are expected to proceed to near completion, according to Fig. 5.

The measured time-courses (Fig. 6) were nearly linear at high and markedly nonlinear at low substrate concentrations, indicating that substrate stabilized dimer in the course of the enzymatic reaction. Substrate consumption was quite low by the end of the recording time (0.7% and 4.3%, respectively), ruling out substrate depletion as an explanation of the nonlinearity. Also, full-size *dh*PPase, which dissociates much slower (Table 2), generated linear P_i_ accumulation curves under identical conditions (Fig. 6).

**Fig. 6.**
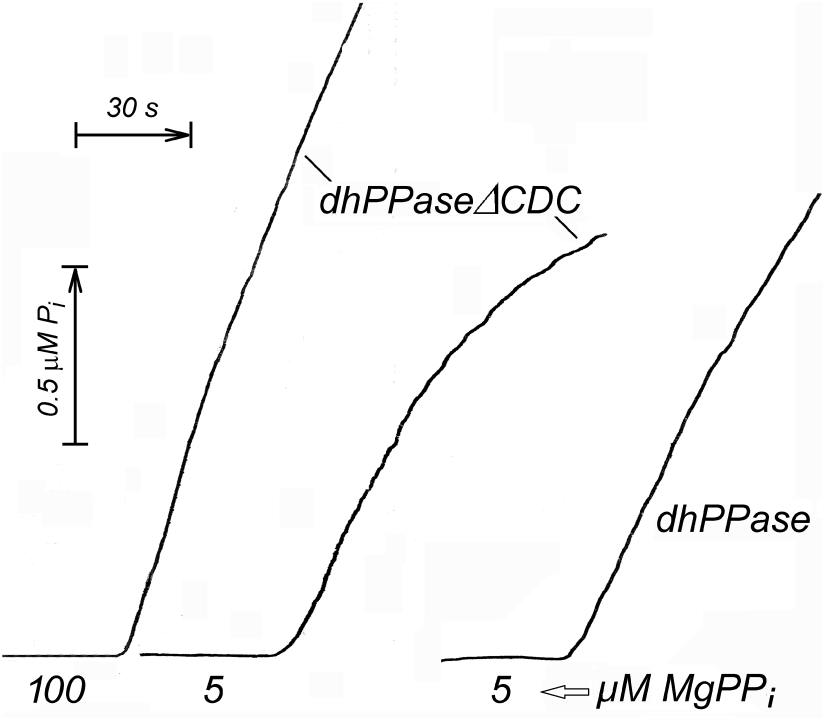
Time-courses of P_i_ production by *dh*PPaseΔCDC and *dh*PPase measured at two substrate concentrations. Total PP_i_ concentration was 280 μM or 14 μM, corresponding to 100 and 5 μM MgPP_i_, as indicated below the curves. Final enzyme concentrations were 0.5 nM (leftmost curve) or 1 nM (two other curves). Stock enzyme solution contained 5.8 µM *dh*PPaseΔCDC or 1 µM *dh*PPase in the isolation buffer; the enzymes were by 37 and 79 % dimeric or tetrameric, respectively, in the stock solution, according to Fig. 5. The phosphate analyzer was operated in a “short dead-time” mode (see Materials and methods) at a sensitivity of 5 μM P_i_ per recorder scale.

Fitting Equation 3 to the time-courses in Fig. 6 yielded inactivation rate constant (*k*_i_) of 1.53 and 0.28 min^-1^ for the *dh*PPaseΔCDC-catalyzed reaction at 5 and 100 µM MgPP_i_, respectively. The former *k*_i_ value is consistent with the *k*_d_ value for *dh*PPaseΔCDC dissociation estimated in the absence of substrate (144 h^-1^, i.e. 2.4 min^-1^; Table 2), keeping in mind that 5 µM substrate partially protected the tetramer against dissociation, i.e., decreased *k*_i_. Based on the above values of the rate constant for dimer dissociation, one can conclude that bound substrate decelerated the reaction by approximately tenfold.

## 4. Discussion

The results reported above indicate that CBS-PPases of five bacterial species are active as homotetramers. Three of these PPases contain the full set of domains (DHH, DHHA2, DRTGG, and two copies of CBS), and the remaining two lack the DRTGG domain. Both CBS-PPase types are regulated by monoadenosine phosphates (AMP, ADP, and ATP) that bind to CBS domains, but only the former type is activated by diadenosine polyphosphates [14], important cellular alarmons [23]. Most other reported PPases are homohexamers of approximately 20-kDa subunits (prokaryotic Family I), homodimers of 32–70-kDa subunits (eukaryotic Family I, prokaryotic canonical Family II, and cation-transporting membrane PPases), or monomers (Family III). Formation of tetramers of 21–25-kDa subunits has been claimed in reports of a preliminary nature on Family I PPases from thermophilic and acidophilic bacteria [24–26]. Conclusive evidence for homotetramer formation has been only reported for a unique Family I vacuolar PPase from *Trypanosoma brucei*, which has an additional EF hand domain in its 47-kDa subunit [27]. CBS-PPase is thus unique among different PPases by the type and size of its oligomer structure.

The first evidence for the tetrameric structure of CBS-PPases came from cross-linking experiments, which revealed tetrameric species after reaction of three CBS-PPases with glutaraldehyde. Interestingly, the cross-linking reaction using glutaraldehyde produced spurious results with the DRTGG-lacking CBS-PPases (*eh*PPase and *el*PPase), and another cross-linker, dimethylsuberimidate, produced only dimers. Notably in this regard, the efficiency of a cross-linking reaction is determined by the existence of a pair of suitably located amino groups across the subunit interface in a dimeric protein and at least two such pairs at different interfaces in a tetramer. The lack of such amino group pairs in *eh*PPase and *el*PPase seems to be the most likely explanation of the low yield of their cross-linked tetramers. This conclusion is supported by a comparison of the lysine content and distribution in different CBS-PPases (Table S3). The total number of lysines per subunit is, on the average, 3.4 times larger in the DRTGG domain-containing CBS-PPases than in their DRTGG domain-lacking counterparts. This difference is mainly due to the low lysine content in the CBS domains of the latter CBS-PPases.

Sedimentation experiments were therefore crucial to confirm tetrameric organization of the CBS-PPases, including those that could not be cross-linked. The critical point in this type of analysis was conversion of the sedimentation coefficient into molecular mass. The samples containing EDTA-treated CBS-PPase (Fig. 3D) were most informative in this regard as they contained a dynamic mixture of all three oligomeric forms — tetramer, dimer, and monomer. An additional reference point was provided by the canonical sgPPase that mimics the catalytic part of CBS-PPase, *dh*PPaseΔCDC, because its dimeric structure was confirmed by X-ray crystallography [8].

Dilution of CBS-PPase solution caused reversible dissociation and inactivation of the tetrameric enzyme. A question to answer is whether the dissociation lead to dimers or monomers. We used the simplest model assuming dimer formation, but, given the relatively high point scatter in Fig. 5A, the models involving tetramer ⇄ monomer or tetramer ⇄ dimer ⇄ monomer equilibrium could not be ruled out. The identification of the oligomeric species generated upon dilution of the full-size enzyme thus requires further investigations.

Reversible dissociation inactivates CBS-PPase and might therefore contribute to regulation of its activity in cells. However, this regulatory mechanism is operable only if enzyme concentration is sufficiently low to allow oligomer dissociation in changing conditions. CBS-PPase concentration in a cell can be estimated based on the rate of PP_i_ production in bacteria. The upper limit for this rate was estimated to be 82.5 mM/min [1], which can be counterbalanced by 3 µM PPase with a turnover number of 450 s^-1^, characteristic of CBS-PPase activated by ATP. At this concentration, CBS-PPase is dissociated by approximately 10% (Fig. 5) and this percentage increases with cellular PP_i_ concentration because PP_i_ stabilizes the tetramer (Fig. 6). However, this mechanism of activity regulation would be far less efficient compared to the mechanism based on adenine nucleotide binding [11, 13].

As the published model of *cp*PPase dimer [12] cannot explain tetramer formation, we have produced alternative models using a Modeller program. The following considerations were taken into account when choosing the principle of tetramer organization. The N- and C-termini of both monomers are located on the same side of the dimeric regulatory insert [12], whereas the two loops within which it is to be inserted are on the opposite sides of the dimeric *bs*PPase [10], which mimics the catalytic part of CBS-PPase. Hence, two dimers of the catalytic part should be arranged as a tetramer in such a way to allow close proximity of two loops belonging to different dimers for fusing with the regulatory domains. Furthermore, two dimers of the catalytic parts should not form a tight contact with each other, and the same refers to the regulatory parts, as none of these parts forms a tetramer separately. In other words, strong contacts that maintain tetramer stability should be formed by catalytic and regulatory parts belonging to different pairs of monomers.

Two modelled structures that best satisfied these requirements were selected. The model that we prefer (Model I) is shown in Fig. 7, and the other one (Model II) is shown in Fig. S4. Both models can be described as a dimer of dimers. Model I positions the regulatory part nearly perpendicular to the plane going through the contact zone of two DHH domains that interact by forming an extended β-sheet both in the modelled structures and in *bs*PPase dimer [10]. The DHH domains of monomers 1 and 2 strongly interact with each other but not with monomers 3 and 4, and vice versa (Fig. 7, left top panel). In contrast, the regulatory parts of monomers 1 and 2 interact crosswise — only with the regulatory parts of monomers 3 and 4, respectively. Accordingly, deletion of either catalytic or regulatory part should prevent tetramer formation. In Model II, each pair of the catalytic domains is rotated by 180^°^ around the axes crossing the tetramer in its largest dimension and by 30° around a perpendicular axis. This allows additional contact of the DHH domains through the α-helices formed by residues 396-409. DHHA2 domains do not participate in oligomeric contacts in Model II, but may form weak contacts in the closed conformation of the catalytic domains in Model I.

**Fig. 7.**
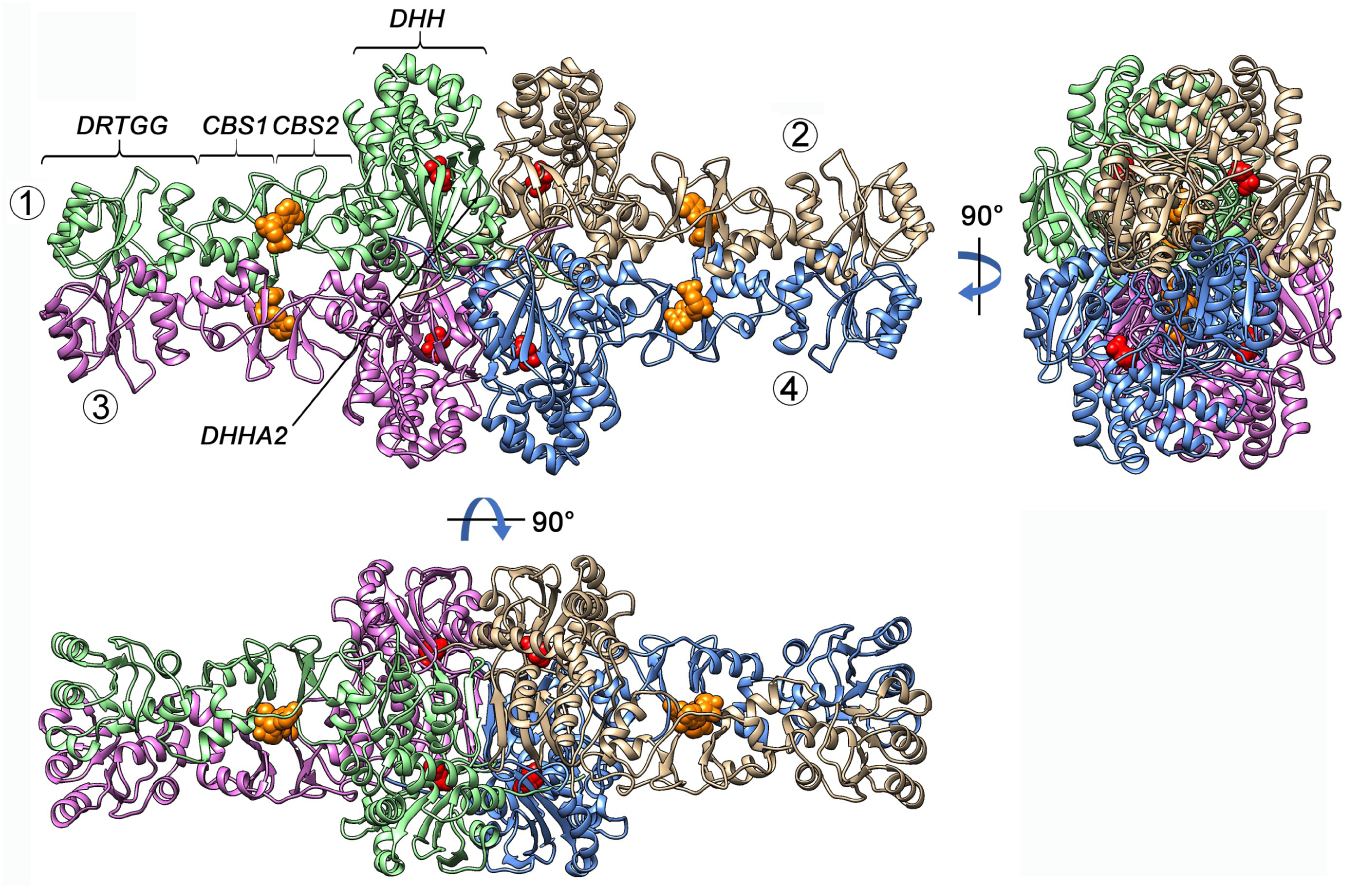
Three views of the modelled *cp*PPase tetramer (Model I). Four monomer units are shown in different colors. AMP bound in the regulatory site and imidodiphosphate bound in the active site are shown as orange and red spheres, respectively. Subunits are numbered and domains are marked in one subunit in the left top panel.

Both models are consistent with known positive co-operativity of CBS-PPase catalysis and regulation. Kinetic co-operativity (decreased *K*_m_ value for interaction with the second substrate molecule) [13] apparently results from interactions in pairs of monomers 1—2 and 3—4, as only this type of interaction occurs in co-operative dimeric Family II PPases devoid of the regulatory CBS domains [28]. AMP, ADP, and ATP are bound by CBS-PPase co-operatively with a stoichiometry of one per monomer [14]. This co-operativity clearly results from interactions within the pairs of the regulatory parts of monomers 1—3 and 2—4, but not between these pairs because they lie far apart in CBS-PPase tetramer. This interpretation is consistent with the non-cooperative binding of adenosine polyphosphates that demonstrated a stoichiometry of one per two monomers [14] because each molecule of this ligand occupies both regulatory sites in each of the two distant monomer pairs [12]. Dilution-induced dissociation of CBS-PPase apparently yields dimers 1—3 and 2—4, which are inactive because the DHH domains cannot form the catalytically competent association via extended β-sheet.

It was of interest to compare CBS-PPase models with published structures of other CBS domain-containing proteins. Most of them form homodimers, with the CBS domains participating in subunit interactions [29], but higher order structures have been also reported. Human cystathionine β-synthase (CBS) contains two catalytic and two CBS domains and appears to be the closest structural analog of CBS-PPase. CBS is homotetrameric in solution. Its structure predicted from determined structure of a dimeric mutant form [30] is a dimer of dimers, wherein strong interactions between catalytic domains in dimer are buttressed by CBS domain-mediated interactions within and between dimers. Like with CBS-PPase, removal of the CBS domains renders CBS dimeric [31]. In homotetrameric putative acetoin dehydrogenase TTHA0829 from *Thermus thermophilus*, the constituting domains have opposite roles, compared to the CBS-PPase models. Its core domain (aspartate-kinase chorismate-mutase tyrA), together with CBS2 domain, participates only in dimer formation, like the CBS and DRTGG domains of CBS-PPase model, whereas two dimers interact only through CBS1 domains that have α-helical and loop extensions [32]. Homooctameric inosine monophosphate dehydrogenase is a dimer of plane tetramers and exists in two forms with different contacts between the tetramers — either predominantly between the catalytic domains or predominantly between the regulatory CBS domains [33]. In the latter case, pairs of the CBS domains (Bateman modules) belonging to different tetramers stack in an unusual antiparallel mode. In tetrameric inosine monophosphate dehydrogenase, the CBS domains do not participate in oligomerization [34]. These structures emphasize variable roles of CBS domain in protein oligomerization.

Further structural studies of CBS-PPase are clearly needed to test the validity of our structural models and provide insight into its regulatory mechanism, which apparently depends on interactions between domains and subunits. Crystallization attempts with CBS-PPase have met with serious difficulties, presumably associated with enormous flexibility of its molecule, the problem common to all CBS domain-containing proteins. Perhaps, different approaches should be accessed in future investigations towards the three-dimensional structure of CBS-PPase.

## Acknowledgment

This work was supported by the Russian Science Foundation (research project 19-14-00063).

